# A focus on details? Inconsistent results on auditory and visual local-to-global processing in absolute pitch musicians

**DOI:** 10.1101/455907

**Authors:** T. Wenhart, E. Altenmüller

## Abstract

Absolute pitch, the ability to name or produce a musical tone without a reference, is a rare ability which is often related to early musical training and genetic components. However, it remains a matter of debate why absolute pitch is relatively common in autism spectrum disorders and why absolute pitch possessors exhibit higher autistic traits. By definition absolute pitch (which involves the analysis of single tones) is characterized by a focus on a local scale than relative pitch (involving relations between tones, intervals, melodies).

This study investigated whether a detail-oriented cognitive style, a concept borrowed from the autism literature (weak central coherence theory), might provide a framework to explain this joint occurrence. Two local-to-global experiments in vision (hierarchically constructed letters) and audition (hierarchically constructed melodies) as well as a pitch adjustment test measuring absolute pitch proficiency were conducted in 31 absolute pitch and 33 relative pitch professional musicians. Analyses revealed inconsistent group differences among reaction time, accuracy and speed-accuracy-composite-scores of experimental conditions (local vs. global, and congruent vs. incongruent stimuli). Furthermore, amounts of interference of global form on judgements of local elements and vice versa were calculated. Interestingly, reduced global-to-local interference in audition was associated with greater absolute pitch ability and in vision with higher autistic traits. Results are partially in line with the idea of a detail-oriented cognitive style in absolute pitch musicians. The inconsistency of the results might be due to limitations of global-to-local paradigms in measuring cognitive style and due to heterogeneity of absolute pitch possessors. In summary, this study provides further evidence for a multifaceted pattern of various and potentially interacting factors on the acquisition of absolute pitch.

## 1 Introduction

Absolute pitch, the ability to name or produce a musical tone without any reference (Takeuchi & Hulse, 1993; Ward, 1999), has frequently been associated with autism (e.g. Bonnel et al., 2003; Heaton, Hermelin, & Pring, 1998; for a review see Mottron et al., 2012) and autistic traits (Brown et al., 2003; Dohn, Garza-Villarreal, Heaton, & Vuust, 2012). Ever since the association was first observed, a potential common framework for both phenomena and possible reasons for their joint occurrence have been matters of debate. Absolute pitch is a rare condition (<1% in the general population, Profita, Bidder, Optiz, & Reynolds, 1988) with a much higher incidence in professional musicians (up to 23%, Deutsch, Henthorn, Marvin, & Xu, 2006; Peter K. Gregersen, Kowalsky, Kohn, & Marvin, 1999, 2001) and people with autism spectrum disorder (e.g. Heaton et al., 1998; Heaton, Williams, Cummins, & Happé, 2008; for a review see Mottron et al., 2012). The latter is defined as a neurodevelopmental condition characterized by difficulties in social verbal and non-verbal communication, and by repetitive behaviors, restricted interests and sensory hyper- or hyposensitivities (Lai, Lombardo, Chakrabarti, & Baron-Cohen, 2013). Several authors have tried to explain absolute pitch with respect to autistic symptoms using theoretical concepts that describe a cognitive style with a tendency towards details. In autism literature, the weak central coherence account (Happé, 1999; Happé & Frith, 2006), the enhanced-perceptional functioning theory (Mottron, Dawson, & Soulieres, 2009; Laurent Mottron, Dawson, Soulières, Hubert, & Burack, 2006), empathizing-systemizing-theory (Baron-Cohen, 2005; Baron-Cohen, 2009) and the theory of veridical mapping (Mottron et al., 2012) are important frameworks, that include the concept of a detail-focused cognitive style. At the same time, Chin (2003) has proposed that absolute pitch musicians may also share a tendency to focus on details, and that this may be associated with an early start in musical training before the age of seven. Chin (2003) argues that the more detailed view of the world that children exhibit up to the age of six (Poirel, Mellet, Houdé, & Pineau, 2008; Poirel et al., 2011) leads to absolute pitch often only developing (or being maintained) during that period. In general, absolute pitch seems to be an excellent model to investigate the interaction of genetic and environmental influences on the acquisition and development of expert abilities (Zatorre, 2003). A large body of research exists on the heritability of absolute pitch (Athos et al., 2007; Baharloo, Johnston, Service, Gitschier, & Freimer, 1998; Peter K. Gregersen et al., 1999), the importance of early musical training and sensitive periods (Deutsch et al., 2006; Peter K. Gregersen et al., 2001; Russo, Windell, & Cuddy, 2003; Schellenberg & Trehub, 2003) and neurophysiological and neuroanatomical differences related to absolute pitch (for a review see Bermudez & Zatorre, 2009; Zatorre, 2003). While some of the neuroscientific results have been discussed against the background of a possible relation between absolute pitch and autism (Dohn, Garza-Villarreal, Chakravarty, Hansen, Lerch & Vuust, 2015; Jäncke, Langer & Hänggi, 2012; Loui, Li, Hohmann & Schlaug, 2011; Loui, Zamm & Schlaug, 2012a), cognitive style (in the sense of a focus on details) in absolute pitch musicians - as compared to the weak-perceptual-coherence account of autism - has never been investigated before.

Typically, paradigms to investigate detail vs. context-based cognition in autism follow the approach of the classical psychophysical experiments by Navon (1977) and consist of hierarchically organized visual (see e.g. Bölte, Holtmann, Poustka, Scheurich, & Schmidt, 2007; Happé, 1999; Happé & Frith, 2006; Mottron, Burack, Iarocci, Belleville, & Enns, 2003) or auditory (e.g. Bouvet, Simard-Meilleur, Paignon, Mottron, & Donnadieu, 2014; Foxton et al., 2003; Justus & List, 2005; List, Justus, Robertson, & Bentin, 2007; Mottron, Peretz, & Menard, 2000) stimuli, e.g. a global letter shape consisting of small letters of either the same or another letter. A range of prior studies have provided evidence for a detail-oriented cognitive style in autistic people in vision (e.g. Bölte et al., 2007; Grice et al., 2001; Mottron et al., 2003; Pring, Ryder, Crane, & Hermelin, 2010; Russell-Smith, Maybery, Bayliss, & Sng, 2012). Recently, Bouvet et al. (2011) developed a paradigm to parallel the experiment in audition. Subjects had to rate the direction of short hierarchically-constructed melodies, where either the whole melody or parts of it were rising or falling. Again, people with autism spectrum disorders showed a detailed-oriented style in this auditory experiment on cognitive style (Bouvet et al., 2014).

If people with autism exhibit a more detail-oriented cognitive style, not only in vision but also in audition, this could be a possible reason for the high frequency of absolute pitch in autistic people, as absolute pitch – by definition - requires focus on a lower-level perceptual entity (single tones), while relative pitch has a broader attentional focus (intervals and melodies, i.e. relation between two or more pitches). However, it is unclear whether healthy absolute pitch possessors show a similar focus on details in vision and or audition, which could explain higher scores on autism self-rating scales. Prior studies have only investigated visuo-spatial abilities (Costa-Giomi, Gilmour, Siddell, & Lefebvre, 2006) and auditory digit span in AP (Deutsch & Dooley, 2013) as well as the relation between relative and absolute pitch abilities in the same subjects (Ziv & Radin, 2014).

To our knowledge, this is the first study to investigate cognitive style in professional musicians with absolute vs. relative pitch, and its relation to accuracy of AP and autistic traits within the same sample. This study will therefore shed new light on the debate on why absolute pitch and autism are frequently associated and whether cognitive style could be their common framework.

## 2 Methods

### 2.1 Setting

The study was part of a larger project consisting of several experiments at the Institute of Music Physiology and Musicians Medicine of the University for Music, Drama and Media, Hannover. Two further experiments and EEG recordings were conducted within the same two sessions in the lab and are reported elsewhere (Wenhart, Bethlehem, Baron-Cohen & Altenmüller, *under review*). For this reason, pitch adjustment assessment as well as cognitive tests from previous publications were also used as control variables here. Therefore, all subjects participated in three parts: an online survey and two appointments in the lab. The online survey was used for pitch identification screening and diagnostic as well as demographic questionnaires (see below). General intelligence tests, a musical ability test, a pitch adjustment test (Anders Dohn, Garza-Villarreal, Ribe, Wallentin, & Vuust, 2014) and two experiments assessing local-to-global processing both in vision and audition were conducted in the lab (see Table 1).

**Table 1.** Participants’ characteristics. Age, nonverbal IQ (SPM), information processing capacity (ZVT), musical training (total hours during life span on main instrument), musicality (AMMA; MSI) and online pitch identification screening (PIS) for each group; * two RP reported not having absolute pitch but reached a screening score of 13 respectively 21. Because of this and their weak performance in the pitch adjustment test, the subjects were assigned to the RP group; Significant group differences are highlighted in bold. AQ refers to autism spectrum quotient (Simon Baron-Cohen, Wheelwright, Skinner, Martin, & Clubley, 2001), MAD = Mean absolute derivation from standard tone, SDfoM = Standard deviation from own mean deviation.

|  | AP (n=31) |  |  | RP (n=33) |  |  | t-test |
| --- | --- | --- | --- | --- | --- | --- | --- |
|  | Mean | SD | Range | Mean | SD | Range |  |
| age | 25.13 | 9.2 | 17-58 | 24.0 | 7.02 | 17-57 | t(56.1)=<br>-0.549;<br>p = 0.585 |
| SPM-IQ | 110.4 | 16.4 | 73-132.25 | 114.41 | 13.14 | 86.5-134.5 | t(57.5)= 1.073;<br>p = 0.288 |
| ZVT-IQ | 120.76 | 13.14 | 101.5-145 | 120.61 | 13.69 | 97-143.5 | t(61.9)=<br>-0.045;<br>p = 0.964 |
| hours main<br>instrument | 11961.4 | 9212 | 1642.5-39785 | 13735.61 | 17125.89 | 1606-77617.25 | t(49.7)= 0.520;<br>p = 0.605 |
| AMMA | 64.74 | 6.26 | 53-78 | 63.244 | 7.03 | 46-76 | t(61.8)= -0.90;<br>p = 0.370 |
| MSI | 208.65 | 17.59 | 161-234 | 210.79 | 15.12 | 185-246 | t(59.3)= 0.521;<br>p = 0.604 |
| **PIS** | 28.5 | 6.03 | 15-36 | 5.30 | 4.33 | 0-21* | **t(52.2)=**<br>**-17.37;**<br>**p < 2.2e-16** |
| **AQ** | 20.48 | 6.05 | 10-36 | 16.88 | 5.44 | 6-27 | **t(60.3)=**<br>**-2.501;**<br>**p = 0.015** |
| **MAD** | 41.37 | 36.49 | 9.8 -200.57 | 296.84 | 86.12 | 91.04 -467.52 | **t(43.7)=**<br>**15.614;**<br>**p < 2.2e-16** |
| **SDfoM** | 52.31 | 44.96 | 7.41-235.69 | 329.77 | 122.77 | 134.37 -811.73 | **t(40.9)=**<br>**12.145;**<br>**p = 3.788e-15** |
| starting age | 5.97 | 2.97 | 2-17 | 7.12 | 2.22 | 3-12 | t(55.4)= 1.751;<br>p = 0.086 |

### 2.2 Participants

In total, 31 AP musicians (16 female) and 33 RP musicians (15 female) participated in the study. The above-mentioned online survey (UNIPARK software, https://www.unipark.com/) was used to recruit participants. They primarily were students or professional musicians at the University for Music, Drama and Media, Hanover; four AP and two RP were amateur musicians. As part of the online survey, a pitch identification screening test (PIS), consisting of 36 categorical, equal-tempered sine tones over a three octave range between C4 (261.63 Hz) and B6 (1975.5 Hz) was used to allocate the participants to groups (AP: >12/36 tones named correctly, else RP). Non-native German speakers had the choice between a German and an English version of the experiments (four AP subjects). All participants but one reported no regular medication or drug intake. None of the participants reported any history of severe psychiatric or neurological condition. The AP group consisted of 15 pianists, 9 string players, 3 woodwind instrumentalists, two singers and 2 brass players; the RP group consisted of 13 pianists, 4 string players, 6 woodwind instrumentalists, 3 bassists/guitarists/accordionists, 3 singers, one drummer and 3 brass players. The Edinburgh Handedness Inventory (Oldfield, 1971) was used to assess handedness. Apart from one subject all AP were consistently right handed, whereas three RP were left-handed and two RP ambidextrous. This study was approved by the local Ethics Committee at the Medical University Hannover. All participants gave written consent.

Two standardized tests were used to assess general nonverbal intelligence and information processing speed: Raven’s Standard Progressive Matrices (Raven, Raven, & Court, 2004) and “Zahlenverbindungstest“(ZVT; Oswald, 2016). AMMA (Advanced Measures of Music Audiation; Gordon, 1989), Musical-Sophistication Index (GOLD-MSI; Müllensiefen, Gingras, Musil, & Stewart, 2014) and estimated total hours of musical training within life span (internal online questionnaire) served to control for musical ability and musical experience.

### 2.3 Experiments and material

#### 2.3.1 Pitch adjustment test (PAT)

All participants performed two absolute pitch tests to assign them to groups AP or RP (pitch identification screening, online) and to assess the accuracy of absolute pitch under controlled conditions (pitch adjustment test, lab). During the pitch adjustment test (PAT; Dohn et al., 2014) participants have to adjust the frequency of a sine wave with random start frequency (220 - 880 Hz, 1Hz steps) and try to hit a target musical note (letter presented centrally on PC screen, e.g. “F# / Gb”) as precisely as possible without the use of any kind of reference. Tones were presented through sound-isolating Shure 2-Way-In-ear Stereo Earphones (Shure SE425-CL, Shure Distribution GmbH, Eppingen, Germany) and participants were allowed to choose their octave of preference. The full test consisted of 108 target notes, presented in semi-random order in 3 blocks of 36 notes each (3*12 different notes per block) with breaks between the blocks. Online pitch modulation was provided by rotating a USB-Controller (Griffin PowerMate NA16029, Griffin Technology, 6001 Oak Canyon, Irvine, CA, USA). Participants could flexibly switch between rough and fine tuning by either turning the wheel (10 cent resolution) or by pressing it down while turning (1 cent resolution). Subjects were given a maximum of 15 seconds for each trial and could confirm their answer by pressing a button on a Cedrus Response Pad (Response Pad RB-844, Cedrus Corporation, San Pedro, CA 90734, USA) to automatically proceed with the next trial. The final frequency at the time of the button press or at the end of the maximum time given was recorded. In both cases, the Inter Trial Interval (ITI) was set to 3000 ms. EEG was measured during the PAT but will be reported elsewhere. The final/selected frequencies in each trial were compared to the nearest target tone (< 6 semitones/600cent). The mean absolute deviation (MAD (1), (Anders Dohn et al., 2014)) from the target tone is given as:
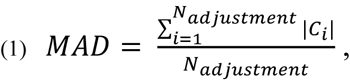

This reflects the pitch adjustment accuracy of the participants and is calculated as the average of the absolute deviations *c*_i_ of the final/selected frequencies from the target tone (referenced to a 440 Hz equal tempered tuning). The consistency of the pitch adjustments (SDfoM, Standard deviation from own mean), possibly reflecting the tuning of the pitch template (Dohn et al., 2014), is then estimated by taking the standard deviation of the absolute deviations (2).
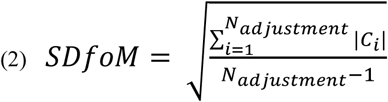

Z-standardization of the MAD (Z_MAD, formula (3)) and SDfoM (Z_SDfoM, formula (4)) values relative to the mean and standard deviation of the non-AP-group were performed for statistical analyses, as originally proposed by Dohn et al. (Anders Dohn et al., 2014).
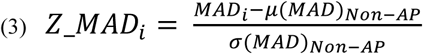

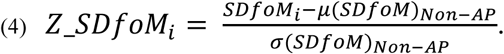

#### 2.3.2 Autistic traits

The Adult Autism Spectrum Quotient (AQ, (Baron-Cohen et al., 2001); German version by C.M. Freiburg, available online: https://www.autismresearchcentre.com/arc_tests) was used to measure autistic traits. The questionnaire was presented within the online survey and consists of 50 items within five subscales (attention to detail, attention switching, imagination, social skills and communication). Items (half of them negatively poled) corresponding to either a mild or strong agreement with the autistic-like symptoms are given one point. The maximum AQ-Score therefore is 50).

#### 2.3.3 Auditory global-local test (AGLT)

Hierarchical melodies were constructed according to Bouvet et al. (2014). Melodies consisted of 9 tones in groups of three (triplets), lasted for 1900ms (210 ms per note) and were presented through sound isolating Shure 2-Way In-ear Stereo Earphones (Shure SE425-CL, Shure Distribution GmbH, Eppingen, Germany). Melodies either successively ascended or descended in steps of two semitones, or the triplets ascended/descended and the next triplet started 6 semitones below respectively above the start of the prior triplet (see Figure 1 (a)). Subjects were asked to judge either the direction of the melody as a whole, or the direction of the triplets in two different blocks of 80 trials each. Compared to Bouvet et al. (2014) we transposed the melodies to 11 different tonalities to avoid that subjects could use absolute pitch cues for the task. One of the transpositions (4 trials, one of each condition) was taken for practice at the beginning of each block. The order of blocks (local vs. global condition) was randomized across subjects and groups. A break was given after the first half (40 trials) of each block. Subjects’ responses were recorded via the Cedrus Response Pad (Response Pad RB-844, Cedrus Corporation, San Pedro, CA 90734, USA), with a right button press for ascending and a left button press for descending. Button colors were randomized across subjects and groups. Reaction times (RT) were calculated relative to the first tone, when a decision at the local respectively global level could be made (local: 2nd note, 210ms; global: 4th note, 630ms) to make reaction times between conditions comparable. During each trial, the word “attention” (German: “Achtung!”) was presented for 1000ms at the center of the screen followed by the sound of the melody (1900ms). Responses were allowed for a further 3100ms after the end of the melody. After this time, or if a button press had occurred, the next trial followed after an ISI of 1000ms.

**Figure 1:**
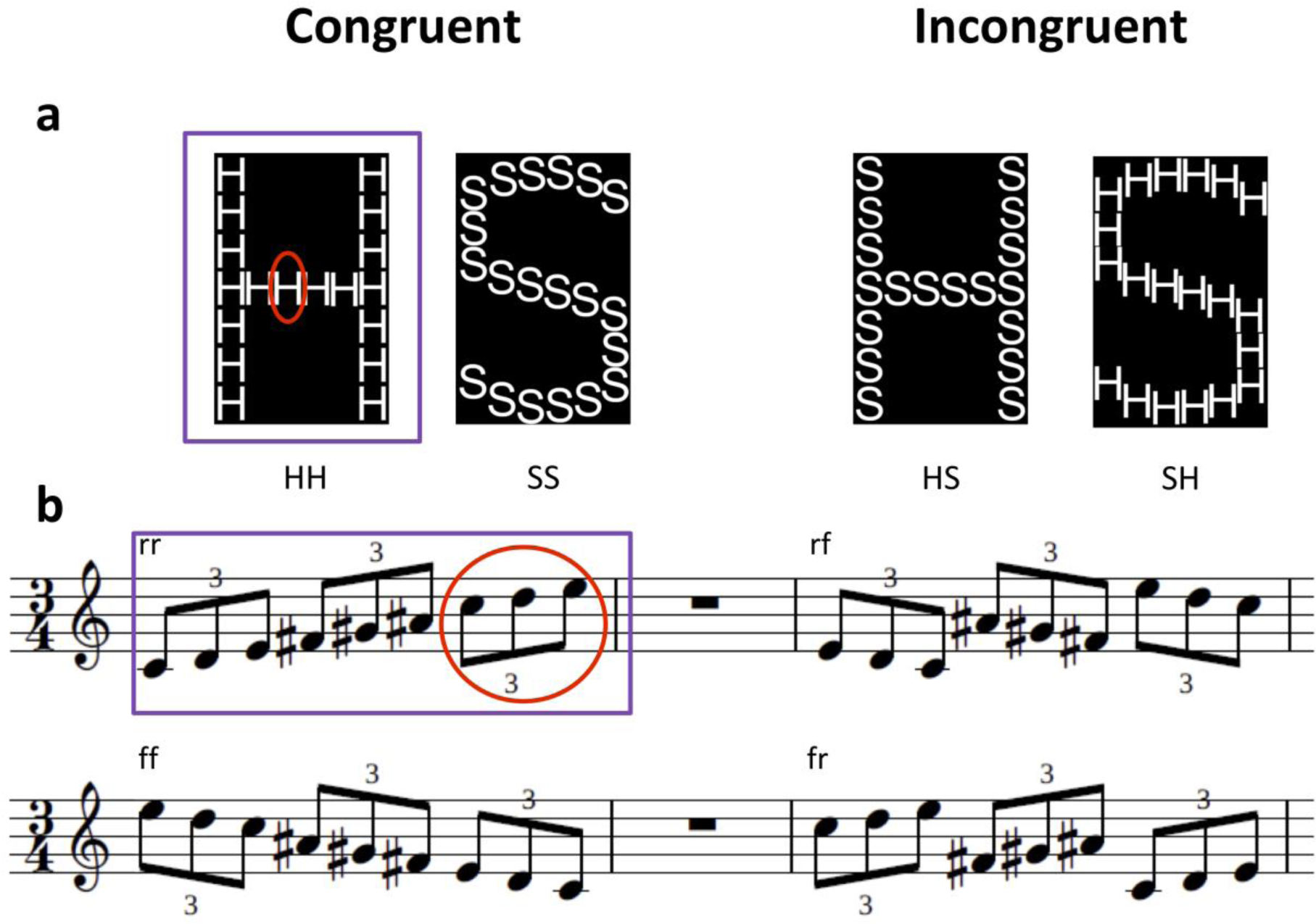
Examples of visual and auditory hierarchical stimuli of the Hierarchical letters (a) and Auditory global-local test (b). Experiments are divided into two blocks, in which participants have to concentrate either on local elements (red; small letters or tone triplets) or on global shape (purple; big letter shape or whole melody). The resulting stimuli can be congruent (e.g. HH: big H, small H; rr= rising tones within whole melody and within triplets) or incongruent (e.g. SH: big S, small H; rf= rising tones within whole melody but falling within triplets). Melodic stimuli occur in different transpositions across all possible tonalities.

A total of nine participants (4 RP, 5 AP) had to be excluded. They either misunderstood the experiment (N=4) or were identified as outliers during inspection of RT distributions (see below) across and within conditions (N=5).

#### 2.3.4 Hierarchical letters (HL)

Four different hierarchical letters were constructed according to Navon (1977). The stimuli were either a global “H” or a global “S” each consisting either of small “H” or small “S” (see Figure 1 (b)). Participants were asked to press a blue button for “H” and a yellow button for “S” or vice versa, depending on randomized allocation of the participants to the two experimental conditions (via the Cedrus Response Pad RB-844, Cedrus Corporation, San Pedro, CA 90734, USA). All participants underwent two blocks of 80 trials each (20 for each stimulus condition). In one block, they were asked to press the two buttons according to the global level of the stimuli, in the other block according to the local level. The order of blocks was randomized across subjects, with half of AP and half of RP starting with the local, respectively global block. Each block had a self-timed break after trial 40. At the beginning of each block, four trials (one per condition) were presented for practice. Within each trial, a fixation cross was present at the center of the screen for 500 ms accompanied by a “beep” sound at the final 75 ms. Afterwards the stimulus was presented for 100 ms in one of the four quadrants around the center of the screen with a visual angle of 4.67° (viewing distance 60 cm; center of the images at [+-2.4,+-2.4] relative to screen center). A dotted mask appeared at the position of the stimulus for 1900ms directly after the end of stimulus presentation, then followed by the next trial. The order of stimuli was randomized and stimulus positions were pseudo-randomized with the same stimulus never occurring twice in a row.

A total of nine participants (3 RP, 6AP) had to be excluded. They either misunderstood the experiment (N=5) or were recognized as outliers during inspection of RT distributions across and within conditions (N=4).

### 2.4 Statistical Analysis

All statistical analyses were conducted using the open-source statistical software package R (Version 3.5 https://www.r-project.org/).

First, only reaction times for correctly answered trials in the experiments HL and AGLT were taken. RT distributions within and across subjects, conditions and groups were inspected. For AGLT, reaction times were calculated relative to the first possible time point of decision, i.e. 2nd note for local trials (RT-210ms) and 4th note for global trials (RT-630ms) to make RT’s comparable between conditions. Trials with physiologically impossible RT’s (i.e. <=0) or extremely long RT’s (>1000ms for HL) were then removed. In a next step, individual outliers defined as exceeding +/-2 times the mean absolute deviation from the median of each subject’s RT distribution were identified and the corresponding trials removed. The remaining trials were considered for further statistical group analysis using median and absolute deviation from median as dependent variables because of non-normality of the subjects’ individual RT distributions (the distribution of RT medians across subjects was normal). The process was performed separately for HL and AGLT trials.

We expected group differences between AP and RP regarding performance on local versus global trials and an interaction between congruency and group for both, local and global trials. Three-way 2x2x2 ANOVAs with two within-subjects factors (congruency, hierarchical level) and one between-subjects factor (group) were performed for each experiment (HL and AGLT) on three dependent variables each: accuracy (ACC), reaction time medians (RT) and a combined score “Speed-accuracy-composite-score” (SACS). The latter has been successfully used by Bouvet et al. (2014) (Bouvet, Simard-Meilleur, et al., 2014) for the auditory global-local paradigm, which served as a template for the present study (AGLT experiment). SACS is calculated as the difference of both scores (ACC (%) and RT), which are z-standardized across all conditions (congruency, hierarchical level) and participants (groups). Therefore, SACS quantifies the performance in one score (e.g. ACC) relative to the other (e.g. RT), so as to deconfound individual strategies - e.g. being fast but not very accurate or being very accurate at the expense of RT. A range of other studies, especially in the field of perception research, have made use of SACS and related scores (Austen & Enns, 2003; Collignon et al., 2010; Glaser, Mendrek, Germain, Lakis, & Lavoie, 2012; Romei, Driver, Schyns, & Thut, 2011). Additionally we performed 2x2 ANOVAs on SACS separately for local and global conditions of HL and AGLT, with between-subjects factor “group” and within-subjects factor “congruency”. To investigate interference effects and their correlation with autistic traits (AQ) and pitch adjustment accuracy (MAD, SDfoM), we calculated individual scores for global-to-local and local-to-global interference for RT, ACC and SACS according to (Bouvet et al., 2011)). Global-to-local interference is calculated as the difference between performance on local congruent minus local incongruent trials, using RT, ACC or SACS. Similarly, local-to-global interference takes global congruent minus global incongruent trials. Both measures reflect the degree of interference of the unattended level (e.g. global) on the rating of the attended level (e.g. local; here: global-to-local interference), which is exhibited for incongruent trials relative to congruent trials. Pearson’s product moment correlations were calculated to estimate the relationship between interference effects and autistic traits respectively absolute pitch performance.

## 3 Results

### 3.1 Auditory processing

Analyses revealed a main effect of hierarchical level for RT, F_RT_(1, 53) = 45.33, p <1.21e-08, 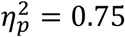, (F_SACS_(1, 53) = 0.17, p = .69; F_ACC_(1, 53) = 1.39, p =.24) and a main effect of congruency for all scores (F_RT_(1, 53) = 34.65, p <2.74e-07, 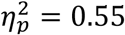; F_SACS_(1, 53) = 33.30, p <4.19e-07, 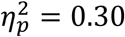; F_ACC_(1, 53) = 36.76, p <1.44e-07, 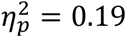). Furthermore, there was a marginally significant main effect of group on RT (F_RT_(1, 53) 3.33, p = 0.07, 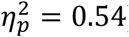). Significant interactions for hierarchical level and congruency (see Figure 2) were also found for all scores (F_RT_(1, 53) = 7.43, p <.009, 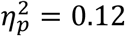; F_SACS_(1, 53) = 25.27, p <6.05e-06), 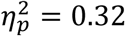; F_ACC_(1, 53) = 23.31, p <1.21e-05, 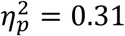 while a significant interaction of group and congruency was only found for ACC (F_ACC_(1, 53) = 4.21, p <.04, 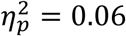) and marginally for SACS (F_SACS_(1, 53) = 3.53, p=0.07, 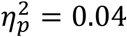). There were no three-way interactions. For means and standard deviations see table 2.

**Figure 2:**
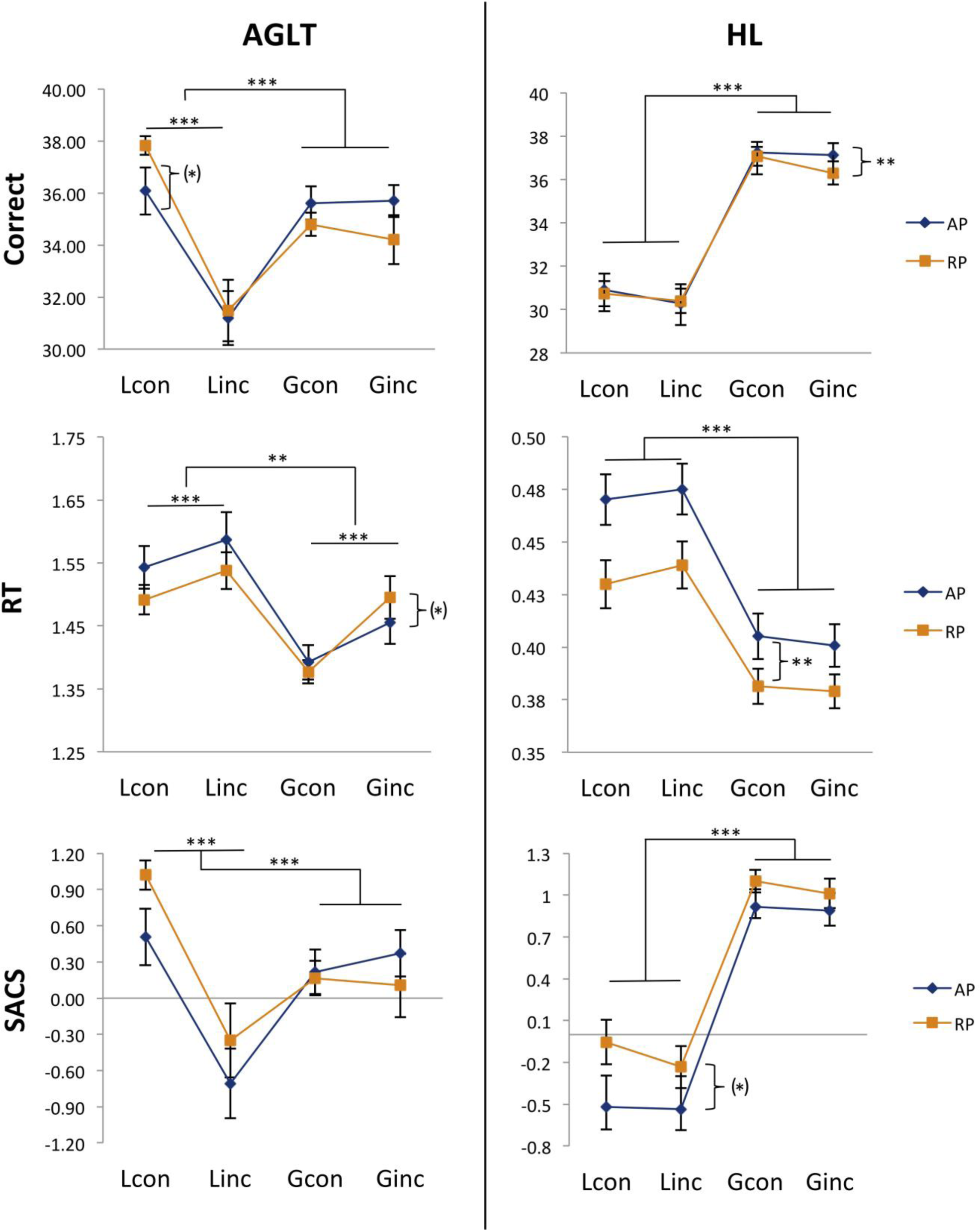
Speed accuracy composite score (SACS), accuracy and RT differences per condition (con= congruent, inc = incongruent; L= local, G= global) and group (AP= absolute pitch, RP= relative pitch). SACS: higher values indicate better performance. Bars represent standard errors. * p<.05. ** p<.01. *** p<.001, (*) p<.10.

**Table 2.** Auditory Processing (N=55) Means (standard deviation) of auditory performance for accuracy (ACC, %), reaction time (RT, ms) and speed-accuracy-composite-score (SACS) by group (absolute pitch, AP, vs. relative pitch, RP).

|  | Global processing |  | Local processing |  |
| --- | --- | --- | --- | --- |
|  | *Congruent* | *Incongruent* | *Congruent* | *Incongruent* |
| **RT** |  |  |  |  |
| AP | 1.39 (0.14) | 1.46 (0.17) | 1.54 (0.17) | 1.59 (0.22) |
| RP | 1.38 (0.10) | 1.49 (0.18) | 1.49 (0.13) | 1.54 (0.16) |
| **ACC** |  |  |  |  |
| AP | 35.61 (3.24) | 35.62 (5.97) | 36.08 (4.65) | 31.19 (5.34) |
| RP | 34.79 (2.37) | 34.79 (3.16) | 37.83 (1.93) | 31.48 (6.39) |
| **SACS** |  |  |  |  |
| AP | 0.22 (0.94) | 0.37 (0.98) | 0.51 (1.60) | -0.71 (2.21) |
| RP | 0.17 (0.80) | 0.10 (1.41) | 1.02 (0.65) | -0.35 (1.65) |

The two-way ANOVA within the local condition revealed a main effect of congruency (F_congruency_(1, 53) = 55.02, p <.9.61e-10, 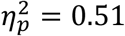) but not of group, nor was there any interaction (F_group_(1, 53) = 1.58, p =0.21, 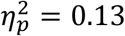; F_congruency x group_ (1, 53) = 1.72, p=0.19, 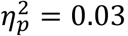). The global condition yielded no significant main effects or interactions (F_group_(1, 53) = 0.39, p =0.53, 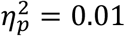; F_congruency_(1, 53) = 0.06, p = 0.81, 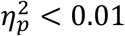; F_congruency x group_ (1, 53) = 1.13, p=0.29, 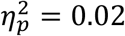). Figure 3 shows differences between conditions per group and experiment.

**Figure 3:**
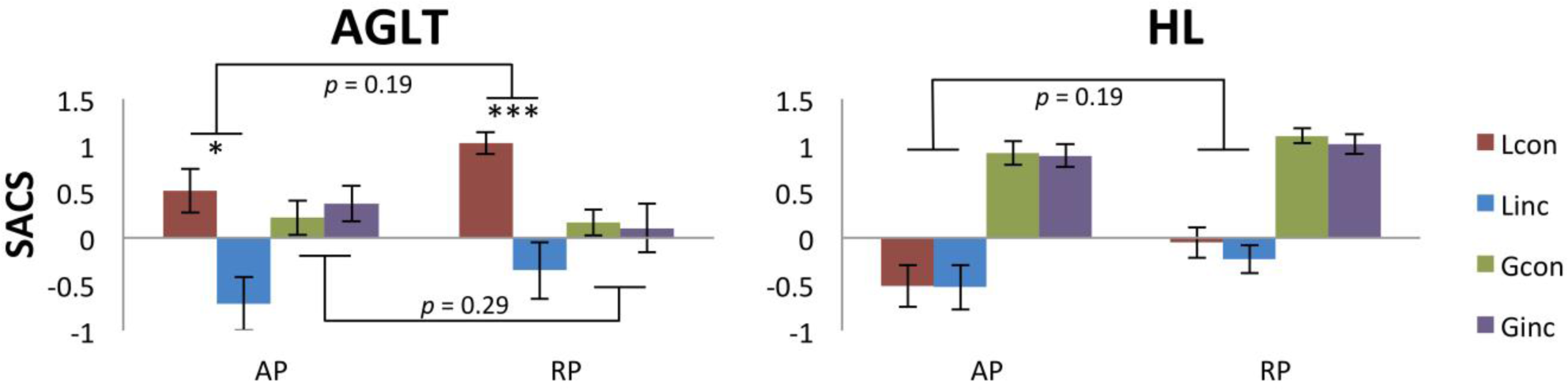
Speed accuracy composite score (SACS) for experimental conditions (hierarchical level, congruency) by group. Left: auditory processing (AGLT), right: visual processing (HL). Marginal significant interaction between group and congruency for AGLT did not reach significance within local vs. global condition. Higher values indicate better performance. HL similarly exhibited a weak tendency for a different effect of congruency within local condition, but remained non-significant. Within-group differences for congruency are shown for all hierarchical levels and both experiments. * p<.05, **p<.01, ***p<.001 (uncorrected)

### 3.2 Visual processing

Analyses yielded main effects of hierarchical level for all scores (F_RT_(1, 53) = 139.19, p <2e-16, 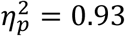, F_SACS_(1, 53) = 94.85, p < 2.1e-13, 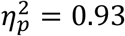; F_ACC_(1, 53) = 119.69, p < 3.31e-15, 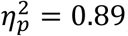) and a marginal main effect of congruency for ACC (F_ACC_(1, 53) = 3.76, p = 0.06, 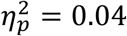). Significant interactions were found for hierarchical level and congruency (see Figure 2) on RT (F_RT_(1, 53) = 5.56, p <.02, 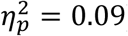), for group and hierarchical level on all scores (F_RT_(1, 53) = 9.58, p <.003, 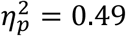; F_SACS_(1, 53) = 4.50, p <0.04, 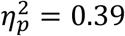; F_ACC_(1, 53) = 6.01, p <.02, 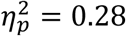), and marginally for group and congruency on RT (F_RT_(1, 53) = 3.86, p = 0.05, 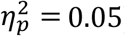). There were no three-way interactions. For means and standard deviations see table 2. Two-way ANOVAs yielded a main effect of group (F_group_(1, 53) = 3.98, p =0.05, 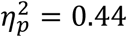) for the local condition (F_congruency_(1, 53) = 1.79, p = 0.19, 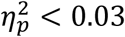; F_congruency x group_ (1, 53) = 0.33, p=0.57, 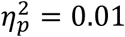) and no effects for the global condition (F_group_(1, 53) = 0.18, p =0.68, 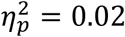; F_congruency_(1, 53) = 0.84, p = 0.36, 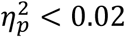; F_congruency x group_ (1, 53) = 0.62, p=0.43, 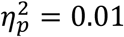).

### 3.3 Interference effects

In general, higher values for local-to-global interference or vice versa indicate higher interference by the local (respectively global) level on incongruent trials. As smaller RT’s indicate better performance, RT interference effects are reversed (lower values indicating higher interference). Analysis of local-to-global interference revealed negative correlations between absolute pitch performance and RT local-to-global interference for the auditory domain (MAD: r=-0.295, p<.05; SDfoM: r=-0.421, p<.001). Therefore higher accuracy on absolute pitch tests (lower values MAD and SDfoM) is associated with weaker local-to-global interference in audition (see Figure 4). No local-to-global interference effects were found for the visual domain.

**Figure 4:**
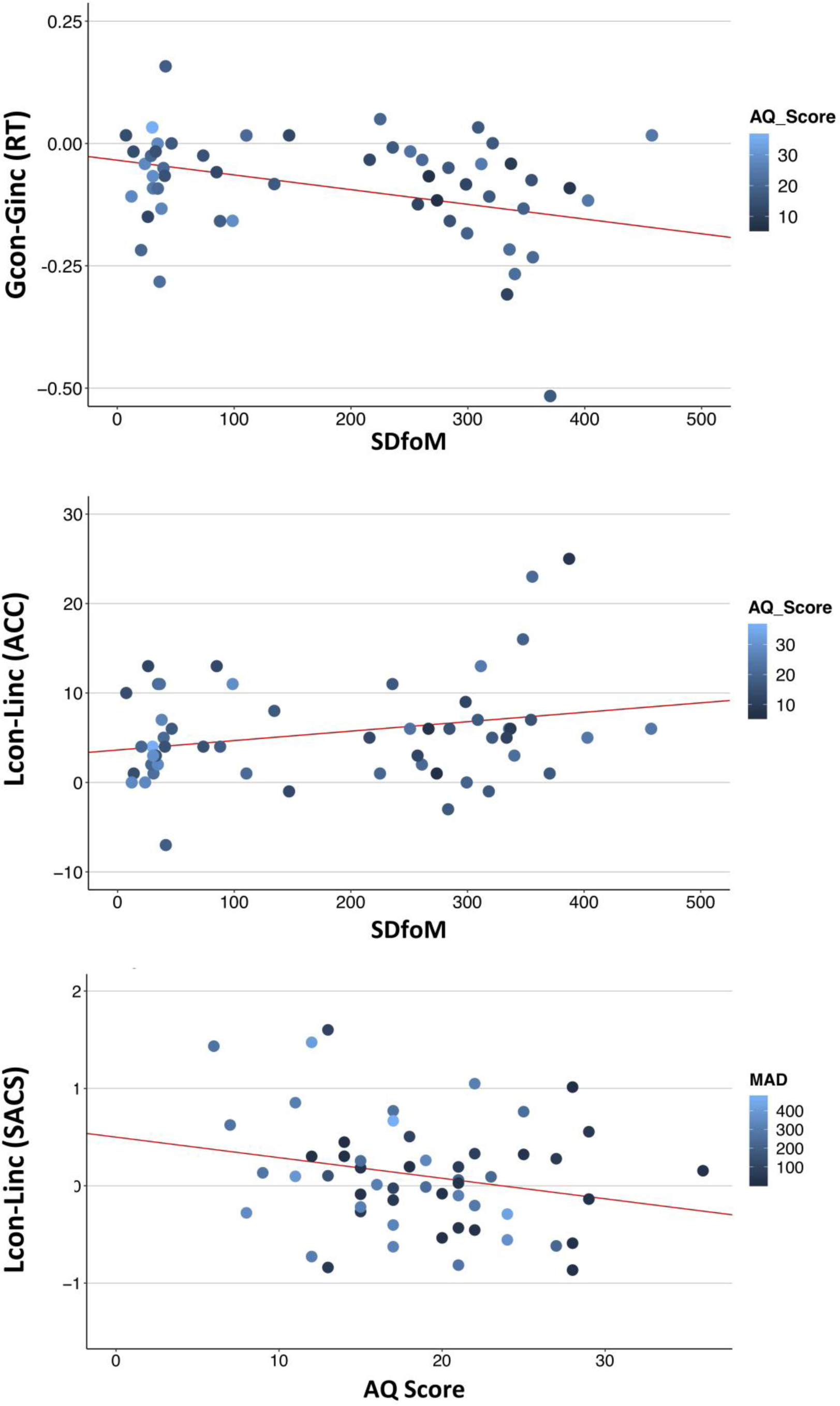
Correlations of visual and auditory interference with autistic traits and absolute pitch performance top: auditory local-to-global interference for RT (reaction times) correlates negatively with standard deviation from target tone in pitch adjustment test; middle: auditory global-to-local interference for ACC (accuracy) correlates positively with standard deviation from target tone; bottom: Visual global-to-local interference (SACS,) correlates negatively with autistic traits (AQ-Score, marginally significant). Higher values for interference (y-axis) indicate higher interference of the first named level (reverse for RT). Colors indicate values for pitch accuracy (MAD) respectively autistic traits (AQ). * p<.05, **p<.01, ***p<.001 (uncorrected).

In the auditory domain, better performance (pitch template tuning, consistency) on absolute pitch tests (SDfoM) was furthermore correlated with reduced global-to-local interference in audition (ACC: r=0.300, p<.05; SACS: r=0.242, p=0.075, marginally significant). Higher autistic traits were associated with marginally lower global-to-local interference in the visual domain (r=-0.231, p=0.090). However, all other correlations remained non-significant (see tables 4 and 5).

**Table 3.**
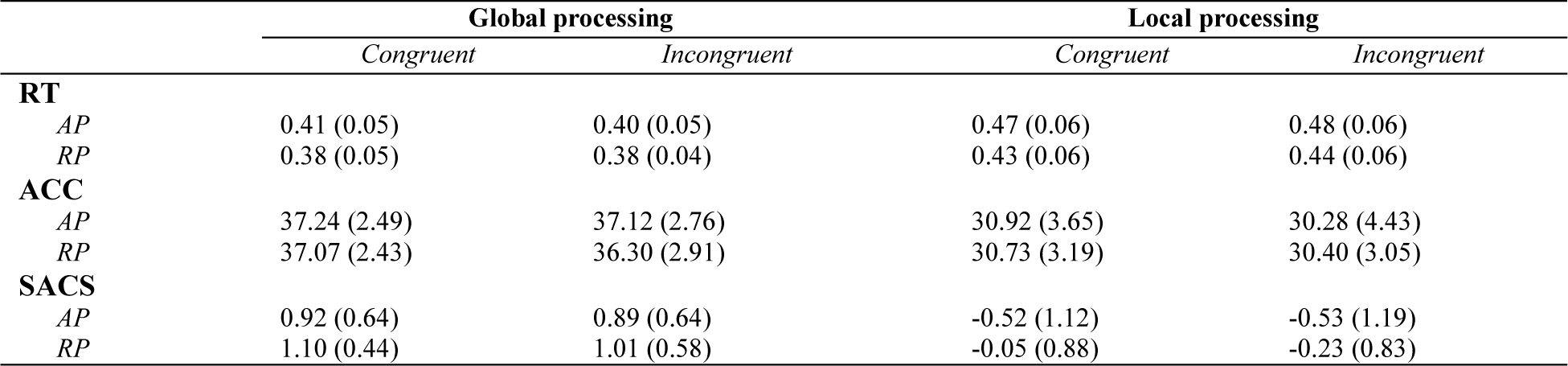
Visual Processing (N=55) Means (standard deviation) of visual performance for accuracy (ACC, %), reaction time (RT, ms) and speed-accuracy-composite-score (SACS) by group (absolute pitch, AP, vs. relative pitch, RP).

**Table 4.** Local-to-global interference (Gcon-Ginc) Pearson’s product moment correlations and p-values for differences between global congruent (Gcon) and global incongruent (Ginc) trials, separately for reaction time (RT, ms), accuracy (ACC, %) and speed-accuracy-composite-score (SACS). Correlations were calculated with autism traits (AQ) and absolute pitch accuracy (MAD, SDfoM).

|  | **AQ** | **MAD** | **SdfoM** |
| --- | --- | --- | --- |
| **RT** |  |  |  |
| AGLT | 0.034 (0.806) | **-0.295 (0.029)** | **-0.421 (<.001)** |
| HL | 0.019 (0.893) | -0.037 (0.789) | -0.033 (0.810) |
| **ACC** |  |  |  |
| AGLT | 0.172 (0.210) | 0.082 (0.550) | 0.096 (0.486) |
| HL | -0.076 (0.583) | -0.024 (0.860) | 0.023 (0.869) |
| **SACS** |  |  |  |
| AGLT | 0.141 (0.304) | 0.105 (0.447) | 0.117 (0.394) |
| HL | -0.038 (0.7845) | -0.064 (0.643) | -0.054 (0.696) |

**Table 5.** Global-to-local interference (Lcon-Linc) Pearson’s product moment correlations and p-values for differences between global congruent (Lcon) and global incongruent (Linc) trials, separately for reaction time (RT, s), accuracy (ACC, %) and speed-accuracy-composite-score (SACS). Correlations were calculated with autism traits (AQ) and absolute pitch accuracy (MAD, SDfoM).

|  | AQ | MAD | SdfoM |
| --- | --- | --- | --- |
| RT |  |  |  |
| AGLT | 0.202 (0.139) | 0.030 (0.827) | -0.006 (0.965) |
| HL | 0.171 (0.213) | -0.013 (0.927) | -0.022 (0.875) |
| ACC |  |  |  |
| AGLT | -0.126 (0.359) | 0.184 (0.178) | **0.300 (0.027)** |
| HL | -0.161 (0.240) | -0.014 (0.919) | -0.114 (0.409) |
| SACS |  |  |  |
| AGLT | -0.171 (0.211) | 0.109 (0.429) | **0.242 (0.075)** |
| HL | **-0.231 (0.090)** | 0.094 (0.495) | 0.018 (0.898) |

## 4 Discussion

The present study is the first to investigate cognitive style, i.e. the tendency to focus more either on details or on the global shape/context of sensory stimuli, in AP and RP musicians. Taken together, our results cannot rule out the hypothesis that AP musicians have a more detail-oriented cognitive style compared to RP, but the evidence is too weak and inconsistent across experiments and conditions, to explain differences in AP performance based on cognitive style alone.

### 4.1 Pitch perception and cognitive style

Performance on auditory or visual hierarchically-constructed stimuli frequently used to assess cognitive style (Bouvet et al., 2011; Navon, 1977) did not reveal strong group differences between AP and RP musicians. As expected, global as well as congruent stimuli revealed a processing advantage both in terms of speed (RT) as well as accuracy (ACC, SACS) independent of group. In general, AP and RP were similar in the degree of performance difference between local and global congruent and incongruent trials (three-way interaction). If anything, the groups might differ in performance on congruent vs. incongruent trials independent of hierarchical level, or vice versa. RT measures in both audition and vision furthermore showed a tendency for slower responses of AP independent of experimental conditions, which was especially prevalent in the visual local condition. Interestingly, groups did not differ in basic information processing speed measured as a confounding variable by ZVT (“Zahlen-Verbindungs-Test”; Oswald, 2016), so general information processing ability cannot account for the differences. In summary, while frequent effects of experimental conditions were independent of groups, there was no consistent tendency towards an advantage for particular processing levels (local vs. global) for the two groups, which would have been reflected in three-way interactions.

Correlation analysis revealed that lower local-to-global interference (higher interference of local percept on global performance) is associated with higher accuracy in pitch adjustment test, but only for RT and only in audition. As we were expecting more detail-oriented perception for AP possessors (Chin, 2003; Mottron et al., 2012), this result actually stands against our hypothesis, as here RP are more affected by details in perceiving a global auditory percept. However, this was only present for RT measures, which alone might not comprise clear evidence in our experiments. Musical stimuli by their nature unfold over time and participants’ response latencies might differ according to their listening strategy. For example some individuals may listen to the whole stimulus, before deciding whether global or local changes were presented, whereas others may choose to press the button as soon as the crucial 4^th^ tone is played (which allows them to notice the difference between global and local stimuli). In line with our hypothesis, reduced global-to-local interference in audition (ACC, SACS) is correlated with higher AP accuracy. In vision however, higher autistic traits are associated with lower global-to-local interference (SACS). Therefore, in audition, people who have a more accurate AP are less affected by the global shape when concentrating on local details, as are people with more autistic traits (in the same sample) in vision. However, we have to admit that this is a weak relationship as it is selective for certain performance measures and sensory domains. In contrast, prior research has shown that cognitive style is quite similar within subjects across audition and vision (Bouvet et al., 2011; Justus & List, 2005; Sanders & Poeppel, 2007). A possible explanation could be that our sample only consists of professional musicians and students at music universities. This is a highly auditorily trained population, a fact which might increase the likelihood of obtaining differing effects in audition and vision as well as potential ceiling effects in audition. Further limitations of our study are the absence of a non-musical control group as well as of a direct comparison to an autistic sample. In general, inconsistent and weak effects might also be due to subgroups within AP musicians, whereby not all AP musicians might exhibit heightened autistic traits and/or a detailed cognitive style. This view receives support from a range of research on AP showing various influences on the acquisition of the ability, including genetics (Baharloo et al., 1998; P. K. Gregersen et al., 2013; Peter K. Gregersen et al., 1999, 2001), an early start of musical training (Baharloo et al., 1998; Bermudez & Zatorre, 2009; Chin, 2003; Gervain et al., 2013; Peter K. Gregersen et al., 2001), a sensitive period (Saffran, 2003; Saffran & Griepentrog, 2001), musical education method (Peter K. Gregersen et al., 2001) and nationality or mother tongue (Deutsch, Dooley, Henthorn, & Head, 2009; Deutsch et al., 2006). However, larger sample sizes are needed to uncover subgroups in such a heterogeneous population.

### 4.2 Hierarchical stimuli and cognitive style

Despite the popularity of the weak-central-coherence account (Happé, 1999; Happé & Frith, 2006) and similar theories of autism (Simon Baron-Cohen, 2009; Mottron et al., 2012, 2006) as well as of the global-local paradigms (Navon, 1977), a few authors have already raised criticism concerning these hypothetical concepts. First, global-local paradigms in the sense of Navon (Navon, 1977) exhibit a huge variability of results even in healthy populations. For example, results are highly affected by relative size and the number of local elements used to construct hierarchical stimuli (Kimchi & Palmer, 1982). Kimchi (1992) further emphasizes that global-local paradigms using hierarchically constructed stimuli might not even measure the degree of holistic perception, as being holistic (i.e. properties that depend on the interrelations between component parts) is not necessarily the same as involving global precedence (i.e. processing of the higher level preceding that of the lower one). Therefore, not all global-to-local paradigms might be adequate to measure holistic perception in terms of Gestalt principles (Wertheimer, 1925). Furthermore, even evidence on a reduced global precedence effect as a result of a more detail-oriented perception in autism is contradictory (Mottron et al., 2003; Mottron, Burack, Stauder, & Robaey, 1999; Mottron et al., 2000; Ozonoff, Strayer, McMahon, & Filloux, 1994).

### 4.3 Future directions

Future studies should therefore address holistic vs. detailed perception using adapted paradigms (e.g. (Kimchi, 1992; List et al., 2007; Sanders & Poeppel, 2007)) to overcome restrictions of classical global-to-local paradigms (Navon, 1977). Furthermore, a consideration of neurophysiological or – anatomical correlates, especially hemispherical contributions, promises to offer a new contribution to the debate of detail-oriented processing style of AP musicians. Seminal work by Peretz and colleagues (1987, 1990) on patients with unilateral brain lesions (Peretz, 1990) and healthy non-musicians (Peretz & Morais, 1987) has shown a processing bias of local information by the left and global by the right hemisphere. This is especially interesting, as research from both fields, autism and absolute pitch, often reveals hemispherical associations (e.g. Brancucci et al., 2009; A. Dohn et al., 2015; Floris et al., 2016; Hyde, Peretz, & Zatorre, 2008; Keenan et al., 2001; Wengenroth et al., 2014; Wilson, Lusher, Wan, Dudgeon, & Reutens, 2009).

To sum up, the correlation analysis of global-to-local interference effects in particular revealed results in accordance with the hypothesis of a more detailed-oriented cognitive style in AP possessors, which is also associated with autistic traits within our sample. However, the inconsistency of the results – and the dissociation of a correlation of AP accuracy with auditory performance versus autistic traits with visual performance - remains to be understood.

## 7 Declarations

### 7.1 Ethics approval and consent to participate

The study was approved by the ethic committee of the Hanover Medical School (Approval no. 7372, committee’s reference number: DE 9515). All participants gave written consent.

### 7.2 Conflict of Interest

The authors declare that the research was conducted in the absence of any commercial or financial relationships that could be construed as a potential conflict of interest.

### 7.3 Author Contributions

TW designed the study, collected, analysed and interpreted the data. EA contributed to the design of the study and interpretation of the data. All authors read, improved and approved the final manuscript.

### 7.4 Funding

TW receives a PhD scholarship from the German National Academic Foundation; TW declares that the funding body has no influence on design of the study and collection, analysis or interpretation of data and in writing the manuscript.

### 7.5 Acknowledgments

We are grateful to Fynn Lautenschläger for support in data collection, Hannes Schmitz, Pablo Carra and Artur Ehle for programming and technical support, and to Dr. Michael Großbach, Christos Ioannou PhD, and Dr. Daniel Scholz on fruitful discussion on the topic and support in data evaluation strategies.

### 7.6 Data Availability Statement

The datasets generated and/ or analyzed during the current study are not publicly available due to specifications on data availability within the ethics approval. Data are however available from the corresponding author upon reasonable request and with permission of the ethics committee of the Hanover Medical School.

